# Molecular Basis for Variations in the Sensitivity of Pathogenic Rhodopsin Variants to 9-*cis*-Retinal

**DOI:** 10.1101/2022.03.01.482516

**Authors:** Francis J. Roushar, Andrew G. McKee, Charles P. Kuntz, Joseph T. Ortega, Wesley D. Penn, Hope Woods, Laura M. Chamness, Victoria Most, Jens Meiler, Beata Jastrzebska, Jonathan P. Schlebach

## Abstract

Over 100 mutations in the rhodopsin gene have been linked to a spectrum of retinopathies that include retinitis pigmentosa and congenital stationary night blindness. Though most of these variants exhibit a loss of function, the molecular defects caused by these underlying mutations vary considerably. In this work, we utilize deep mutational scanning to quantitatively compare the plasma membrane expression of 123 known pathogenic rhodopsin variants in the presence and absence of the stabilizing cofactor 9-*cis*-retinal. We identify 69 retinopathy variants, including 20 previously uncharacterized variants, that exhibit diminished plasma membrane expression in HEK293T cells. 67 of these apparent class II variants exhibit a measurable increase in expression in the presence of 9-*cis*-retinal. However, the magnitude of the response to this molecule varies considerably across this spectrum of mutations. Evaluation of the observed shifts in relation to thermodynamic estimates for the coupling between binding and folding suggests underlying differences in stability constrains the magnitude of their response to retinal. Nevertheless, estimates from computational modeling suggest many of the least sensitive variants also directly compromise binding. Finally, we evaluate the functional properties of three previous uncharacterized, retinal-sensitive variants (ΔN73, S131P, and R135G) and show that two retain residual function *in vitro*. Together, our results provide a comprehensive experimental characterization of the proteostatic properties of retinopathy variants and their response to retinal.

## Introduction

Mutations in integral membrane proteins are responsible for a variety of genetic diseases.(*1, 2*) Most such mutations generate a loss of function (LOF) as a result of one or more molecular defects ranging from the disruption of protein folding, the attenuation of protein expression, changes in protein localization, and/ or the perturbation of a protein’s intrinsic activity.(*2, 3*) An understanding of the molecular defects caused by specific mutations can provide a decisive advantage in drug discovery and targeting.(*4*) For instance, mechanistic knowledge of the effects of common cystic fibrosis transmembrane conductance regulator (CFTR) variants facilitated the successful development of both corrector compounds that restore the expression of misfolded variants (class II) and potentiator compounds that enhance the gating of inactive variants (class III & IV) that are responsible for cystic fibrosis (CF).(*5, 6*) Though the emergence of clinical sequencing has accelerated the discovery of MP variants that are associated with a variety of diseases, experimental efforts to characterize the molecular effects of their mutations have not kept pace.(*4*) Therefore, new experimental approaches that enable rapid characterization of disease-linked membrane protein variants and how they respond to therapeutic lead compounds are needed to guide the development of precision therapeutics.(*2*)

There are currently over 100 known mutations in the rhodopsin G protein-coupled receptor (GPCR) that cause a spectrum of visual retinopathies including autosomal dominant retinitis pigmentosa (adRP) and congenital stationary night blindness (CSNB).(*7*) Most experimentally characterized adRP variants accumulate in the endoplasmic reticulum (ER) in a manner that compromises their maturation within the secretory *pathway*.(*8–10*) In contrast, CSNB variants are typically well expressed, but exhibit constitutive activation.(*11*) Nevertheless, there are wide variations in the age of onset and severity of the retinopathies that likely reflect differences in the molecular effects of these mutations and other uncharacterized variants.(*7*) The expression of many of retinopathy variants can be partially restored by analogs of rhodopsin’s native 11-*cis*-retinal cofactor and/ or other small molecules that bind and stabilize the opsin apoprotein. (*12–16*) Despite the discovery of numerous therapeutic lead compounds and the initiation of several clinical trials, there are currently no approved treatments for these disorders. An improved understanding of the molecular effects of the spectrum of clinical rhodopsin variants may help to identify a subset of “correctable” rhodopsin variants that could be targeted in future clinical trials.

In the following, we apply deep mutational scanning (DMS) to quantitatively compare the plasma membrane expression (PME) of 123 known rhodopsin variants that are associated with visual retinopathies, including 42 that were previously uncharacterized. We show that 69 of these 123 variants exhibit deficient PME in HEK293T cells, including 20 that were previously uncharacterized. Our results reveal that the mutations that have the most severe proteostatic effects on the opsin apoprotein cluster within the protein core and/ or retinal binding pocket. Of the 69 putative class II variants, 67 exhibit a measurable increase in expression in the presence of 9-*cis*-retinal-a photostable isomer of rhodopsin’s native 11-*cis*-retinal cofactor. Nevertheless, response to retinal varies greatly across this spectrum of variants. A comparison of the observed effects of retinal to theoretical estimates of the stabilization afforded by retinal binding suggests that responses are generally constrained by stability. Nevertheless, binding calculations imply that many of the least responsive variants also directly disrupt retinal binding. Finally, we show that two of the three previously uncharacterized variants that exhibit the largest change in PME in the presence of retinal are capable of regenerating rhodopsin pigments that retain residual signaling activity *in vitro*. Together, our findings provide a comprehensive overview of the proteostatic effects of pathogenic rhodopsin variants that may help to guide the discovery and targeting of rhodopsin corrector molecules.

## Results

### Survey of the Plasma Membrane Expression of Retinopathy Variants

To measure the proteostatic effects of retinopathy mutations by DMS, we first assembled a pooled genetic library of containing 119 adRP variants and four CSNB variants. This group of 123 missense and single-codon deletion variants includes 42 previously uncharacterized variants, 57 known class II variants, and 24 variants with other classifications (classes I, and III-VII, Table S1). These mutations are distributed across the primary structure of rhodopsin (Table S1). To ensure even sampling, we generated a set of individual plasmids in which each variant can be matched to a single unique molecular identifier sequence (UMI). A stoichiometric mixture of these plasmids was then used to create a pool of recombinant HEK293T cells in which each cell inducibly expresses a single retinopathy variant from a defined genomic locus, as was described previously.(*17, 18*) A flow cytometry analysis of opsin variant surface immunostaining reveals that ~51% of these cells express variants with comparable PME to wild-type (WT), while the remaining cells express variants with comparable staining to the class II P23H variant (Figure 1). The bimodal nature of this distribution reflects the fact that some retinopathy variants compromise PME (class II) while others simply perturb signaling (classes I and III-VII).(*7*) The relative proportion of cells expressing P23H-like variants decreases by 6% in the presence of 5 μM 9-*cis*-retinal, which suggests the PME of many class II variants can be partially restored by this investigational corrector.

**Figure 1.**
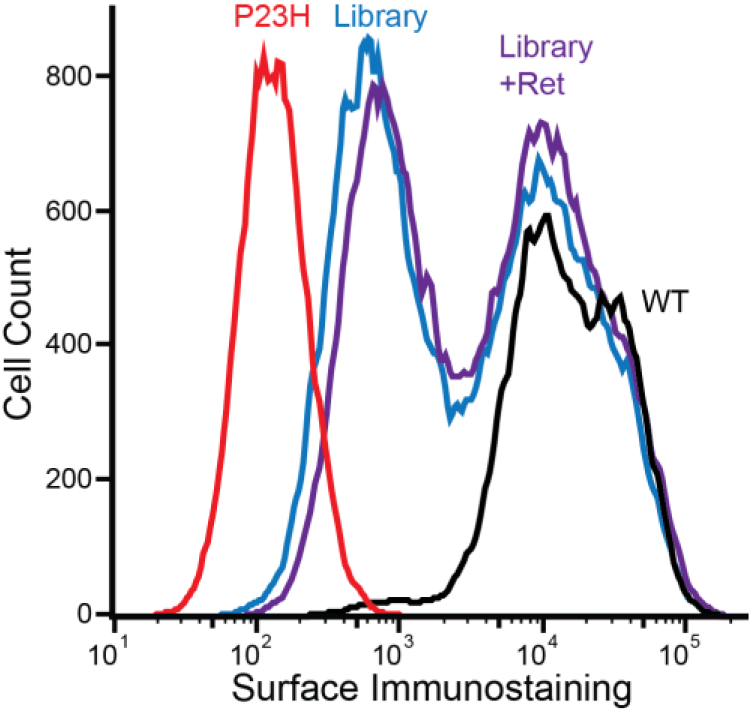
Surface immunostaining profiles of recombinant HEK293T cells expressing rhodopsin variants. A histogram depicts flow cytometry measurements of rhodopsin surface immunostaining intensities among recombinant HEK293T cells that stably express WT opsin (black), P23H opsin (red), or a mixture of retinopathy variants in the presence (purple) and absence (blue) of 5 μM 9-*cis*-retinal.

To estimate the PME of individual variants, we utilized fluorescence activated cell sorting (FACS) to fractionate these cells according to surface immunostaining, extracted the genomic DNA from each fraction, and used deep sequencing of the recombined UMIs to quantify the relative abundance of each variant within each fraction. Sequencing data were then used to estimate the surface immunostaining intensity of each variant, as was described previously.(*18*) Consistent with expectations, known class II variants exhibit lower intensities relative to WT (Average Intensity = 17,400 ± 1,800) and other variants that exhibit other types of conformational defects (Mann-Whitney *p* = 1.9 × 10^−11^, Figure 2A). The distribution of surface immunostaining intensities among previously uncharacterized variants spans the range in between those of previously characterized variants (Figure 2A). This collection of retinopathy variants features a prominent cluster with little to no detectable plasma membrane opsin (severe class II, 44 variants) and a cluster with surface immunostaining intensities that are comparable to WT (other classes, 54 variants, Figure 2B). Nevertheless, there are also several variants with intermediate surface immunostaining intensities (moderate class II, 25 variants, Figure 2B). A projection of variant intensity values onto the structure of rhodopsin reveals that mutations of buried residues near the retinal binding pocket generally cause the largest reduction in PME (Figures 2 A & C). By comparison, mutations within the disordered C-terminal tail have similar expression to WT (Table S1). Variants that are predicted to compromise the translocon-mediated membrane integration of TM domains(*19*) or to disrupt the stability of the native fold generally exhibit attenuated expression (Figure 2A). Together, these results unambiguously identify a comprehensive set of retinopathy variants with attenuated PME and suggest that their proteostatic effects generally arise from perturbations of co- and/ or post-translational folding.

**Figure 2.**
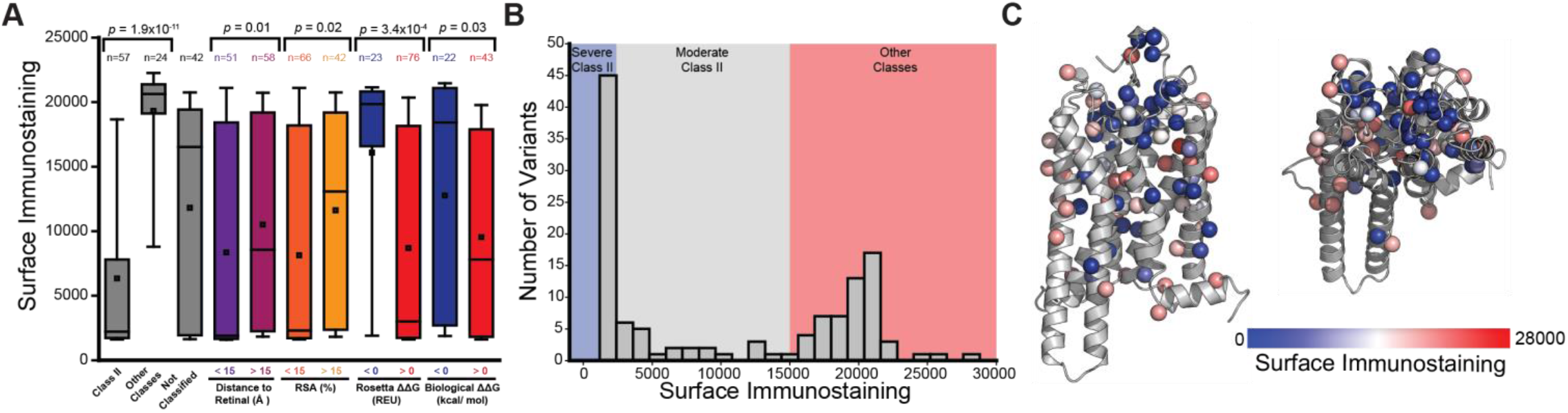
Surface immunostaining intensities of individual opsin variants. Surface immunostaining intensities for individual variants were determined in the absence of retinal by deep mutational scanning (DMS). A) A box and whisker plot depicts the distributions of surface immunostaining intensities among variants that are known to cause misfolding (class II) relative to those known to cause other conformational defects (other classes) and those that are previously uncharacterized (not classified). Distributions are also shown for subsets of variants that are grouped according to whether they occur at positions that are close to the retinal centroid (<15 Å), far from the retinal centroid (>15 Å), buried in the protein core (<15% relative surface area), or solvent exposed (>15% relative surface area). Distributions for mutations that are predicted by Rosetta to stabilize (ΔΔG <0) or destabilize (ΔΔG >0) the native conformation, or that are predicted by the biological hydrophobicity scale to enhance (ΔΔG <0) or disrupt (ΔΔG >0) translocon mediated membrane integration are shown for reference. B) A histogram depicts the range of observed surface immunostaining intensities among the 123 retinopathy variants. Blue, gray, and red regions reflect the intensity intervals corresponding to the designations for severe class II, moderate class II, and other classifications, respectively. C) Intensity values for individual variants are projected onto the corresponding mutated side chains in the three-dimensional structure of rhodopsin (PDB 3CAP). Side chain C_β_ atoms (or glycine H) are colored according to the average intensity from three replicate DMS experiments. Blue indicates poor expression, white indicates intermediated expression, and red indicates expression levels comparable to WT.

### Impacts of 9-cis-retinal on the plasma membrane expression of retinopathy variants

The proteostatic effects of retinal varies considerably across the spectrum of destabilized rhodopsin variants. (*16, 18, 20*) We therefore repeated these experiments in the presence of 5 μM 9-*cis*-retinal to identify retinopathy variants that are most amenable to correction. Consistent with our recent findings,(*20*) the results show that poorly expressed variants exhibit the largest change in expression in the presence of retinal (Figure 3A). There are clusters of highly responsive variants that include several previously uncharacterized mutations near the binding pocket (L47R and T289P) and the cytosolic interface (L57R and ΔN73) (Table S1, Figure 3B). Nevertheless, a structural map of mutated side chains reveals that responsive and non-responsive mutants are found throughout the structure (Figure 3B). This observation implies that, outside of a few select variants, increases in expression are unlikely to arise from structurally-specific interactions between mutated side chains and retinal itself. Our recent findings suggest, instead, that these variations likely reflect the composite effects of mutations on various molecular features that are tied, directly or indirectly, to changes in conformational stability.(*20*) Consistent with this possibility, we find that absolute increases in expression are most pronounced among moderate class II variants (Figure 3C), which perturb different structural regions but generate a comparable reduction in expression. Together, these results identify a subset of “correctable” retinopathy variants and imply that proteostatic responses may be tied to differences in opsin/ rhodopsin stability.

**Figure 3.**
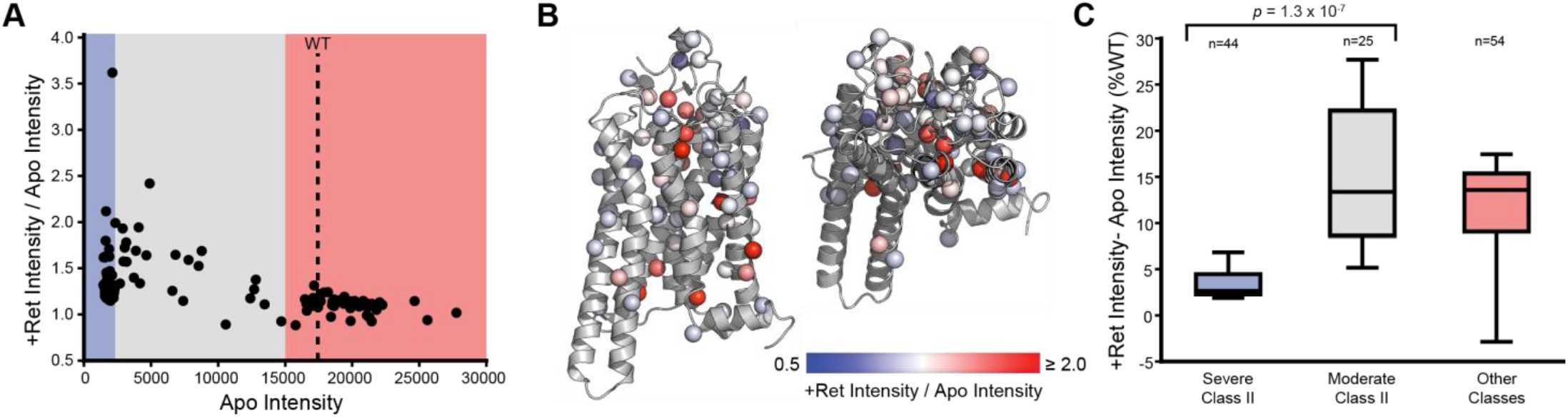
Impact of 9-*cis*-retinal on the surface immunostaining of rhodopsin variants. Surface immunostaining intensities for individual variants are compared in the presence and absence of 5 μM 9-*cis*-retinal by deep mutational scanning (DMS). A) The ratio of the surface immunostaining intensity in the presence of retinal to the intensity in the absence of retinal is plotted against the corresponding surface immunostaining values for each variant in the absence of retinal. Blue, gray, and red regions reflect the intensity intervals corresponding to the designations for severe class II, moderate class II, and other classifications, respectively. B) The ratio of the surface immunostaining intensity in the presence of retinal to the intensity in the absence of retinal for individual variants is projected onto the corresponding mutated side chains in the three-dimensional structure of rhodopsin (PDB 3CAP). Side chain C_β_ atoms (or glycine H) are colored according to the average intensity ratio from three replicate DMS experiments. Blue indicates minimal change in expression, white indicates an intermediate increase in expression, and red indicates a large increase in surface immunostaining intensity in the presence of retinal. C) A box and whisker plot depicts the distribution of the difference between surface immunostaining intensities in the presence and absence of retinal normalized relative to the WT intensity among severe class II (blue), moderate class II (gray), and variants from other classes (red).

### Energetic Interpretation of the Observed Trends in Variant Expression

The sensitivity of moderate class II variants to retinal potentially arises from the energetic coupling between binding and folding. To rationalize the energetic basis of these observed trends, we used a series of simplifying assumptions to approximate the stability of each variant, how much retinal binding should increase variant stabilities, and how much this stabilization should increase variant expression levels (see *Materials and Methods*). These gross simplifications provide a lens to understand how differences in stability shape the response to retinal. First, we note that retinal should have no effect on variants that compromise binding. These variants should therefore fall along a diagonal when rhodopsin variant intensities (+ retinal) are plotted against their corresponding opsin intensities (apo) (red dashes, Figure 4A). In contrast, variants with native binding energetics should exhibit an increase in intensity proportional to the change in the combined fraction of folded opsin and rhodopsin (*f*_fold_), which can be calculated by combining ΔG_fold_ values estimated from the expression of each variant with the estimated free energy of binding (ΔΔG_fold_ ~ 1.1 kcal/ mol) (blue dashes, Figure 4A). The observed changes in variant immunostaining intensities generally fall between these bounds (Figure 4A). While our data lack true internal measurements for WT, estimated intensities from independent measurements(*21*) place the shift in WT expression close to the upper bound (Figure 4A, purple). Consistent with the observed trends (Figure 3C), the general properties of this model and the resulting shape of the upper boundary suggest marginally stable variants (apparent ΔG_fold_ ~ 0 kcal/ mol) should exhibit the largest absolute increases in intensity (blue dashes, Figure 4A, Figure S1). Nevertheless, relatively few variants exhibit gains in expression that approach the upper bound (Figure 4A), which suggests the response to retinal is also likely sensitive to other effects of these mutations.(*20*)

**Figure 4.**
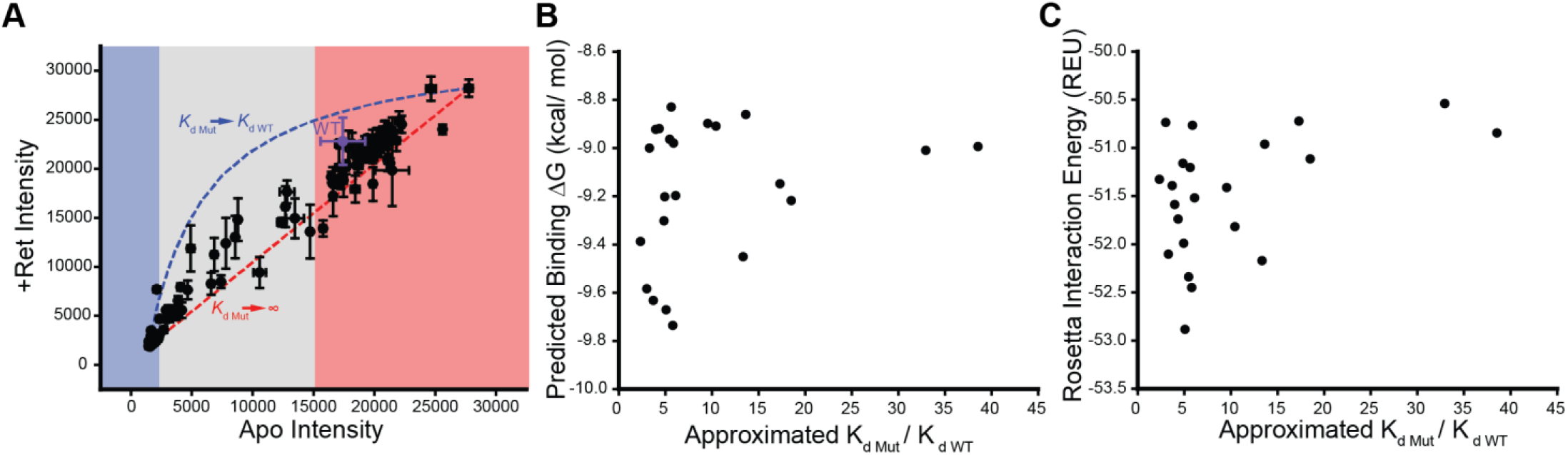
Thermodynamic interpretation of the proteostatic effects of retinal. A) Deep mutational scanning measurements of the surface immunostaining intensities of individual variants in the presence of 5 μM 9-*cis*-retinal (+ Ret Intensity) are plotted against the corresponding values in the absence of retinal (Apo Intensity). Values represent the average of three biological replicates and brackets reflect the standard deviation. Blue, gray, and red regions reflect the intensity intervals corresponding to the designations for severe class II, moderate class II, and other classifications, respectively. The blue dashed line reflects the upper boundary for the increase in intensity for variants that retain WT binding affinity. The red dashed line reflects the lower intensity boundary for variants that do not bind retinal. B) 9-*cis*-retinal was non-covalently docked into structural models of the 25 intermediate class II variants and the corresponding binding energy was predicted using the *K*_DEEP_ web server.(*23*) Predicted binding energies are plotted against thermodynamic approximations for the increase in K_d_ that were calculated based on the observed changes in variant immunostaining intensities. C) Rosetta was used to estimate the protein-ligand interface energy in the context of structural models of the 25 intermediate class II variants featuring a Schiff base between 9-*cis*-retinal and K296. Rosetta interface energies are plotted against thermodynamic approximations for the increase in *K*_d_ that were calculated based on the observed changes in variant immunostaining intensities.

The insensitivity of certain variants to retinal could reflect secondary effects of these mutations on the binding affinity.(*20*) To evaluate whether changes in binding energetics appreciably contribute to variations in the proteostatic effects of retinal, we analyzed the observed expression patterns in relation to computational estimates for the effects of mutations on binding. We first assumed that deviations from the upper bound arise solely from the effects of mutations on the binding energy. Using the framework described above, we estimated the impact of retinal on the f_fold_ and ΔG_fold_ for each variant from observed changes in immunostaining intensity. Based on the change in the apparent ΔG_fold_ (ΔΔG_fold_), we then calculated the corresponding change in the retinal equilibrium dissociation constant relative to WT (*K*_d Mut_ / *K*_d WT_).

To determine whether these projected variations in binding can be reconciled with the structural effects of individual mutations, we carried out two distinct structural analyses. We first utilized Rosetta to generate structural models of both the opsin (apo) and rhodopsin (+ retinal) forms of the target variants. To survey perturbations of the initial binding reaction, we used RosettaLigand(*22*) to dock 9-*cis*-retinal into the binding pocket of each WT opsin and used this docked structure as a template to build structural models of each variant. We then used a convolutional neural network (*K*_DEEP_)(*23*) to predict changes in the free energy of binding. To survey perturbations of the mature pigment, we used Rosetta to calculate the change in the protein-ligand interface energy in the context of rhodopsin variant models bearing the native Schiff base linkage between K296 and 9-*cis*-retinal. Both sets of calculations suggest retinal-sensitive mutants (*K*_d Mut_ / *K*_d WT_ ≤ 10) vary with respect to their predicted effects on the retinal binding energy (Figure 4 B & C). However, both analyses also show that the variants that are least sensitive to retinal (*K*_d Mut_ / *K*_d WT_ ≥ 10) are generally predicted compromise binding (Figures 4 B & C). We should note that we restricted this analysis to the 25 intermediate class II variants due to dynamic range constraints; the difference between the upper and lower bounds approaches the magnitude of experimental variation for variants with the highest or lowest expression (Figure 4A). Thus, it is unclear whether changes in binding energetics are likely to factor into the retinal-sensitivity of this entire spectrum of variants. Nevertheless, these results suggest that mutation-specific responses can be generally reconciled with energetic perturbations of binding and/ or folding equilibria.

### Functional Characterization of Retinal-Sensitive Variants of Uncertain Significance

Though our analyses identify several previously unclassified variants that exhibit enhanced expression in the presence of 9-*cis*-retinal (Table S1), it is unclear whether any of these variants are likely to regain function. To assess the potential functional relevance of these observed proteostatic effects, we characterized the biochemical properties of two previously unclassified intermediate class II variants (ΔN73 and R135G) and one previously unclassified severe class II variant (S131P) that recover expression in the presence of retinal (Table S1). Briefly, we transiently expressed each of these variants in HEK293T cells in the presence of 9-*cis*-retinal, harvested cellular membranes and purified each variant into n-dodecyl-β-D-maltopyranoside (DDM) micelles. Absorbance spectra of each purified variant reveals that both intermediate class II variants (ΔN73 and R135) are at least partially capable of binding retinal and recovering the native absorbance of WT rhodopsin (Figure 5A). Moreover, these variants retain the ability to partially activate G_t_ in response to photoactivation (Figure 5B). In contrast, the severe class II variant S131P failed to regenerate the native pigment even though treatment with this molecule nearly doubles its PME (Figure 5A, Table S1). This observation potentially suggests this proteostatic response arises from cotranslational interactions.(*20*) Together, these findings reveal that certain intermediate class II variants are likely to retain residual activity upon recovery of expression but raise doubts about the druggability of severe class II variants.

**Figure 5.**
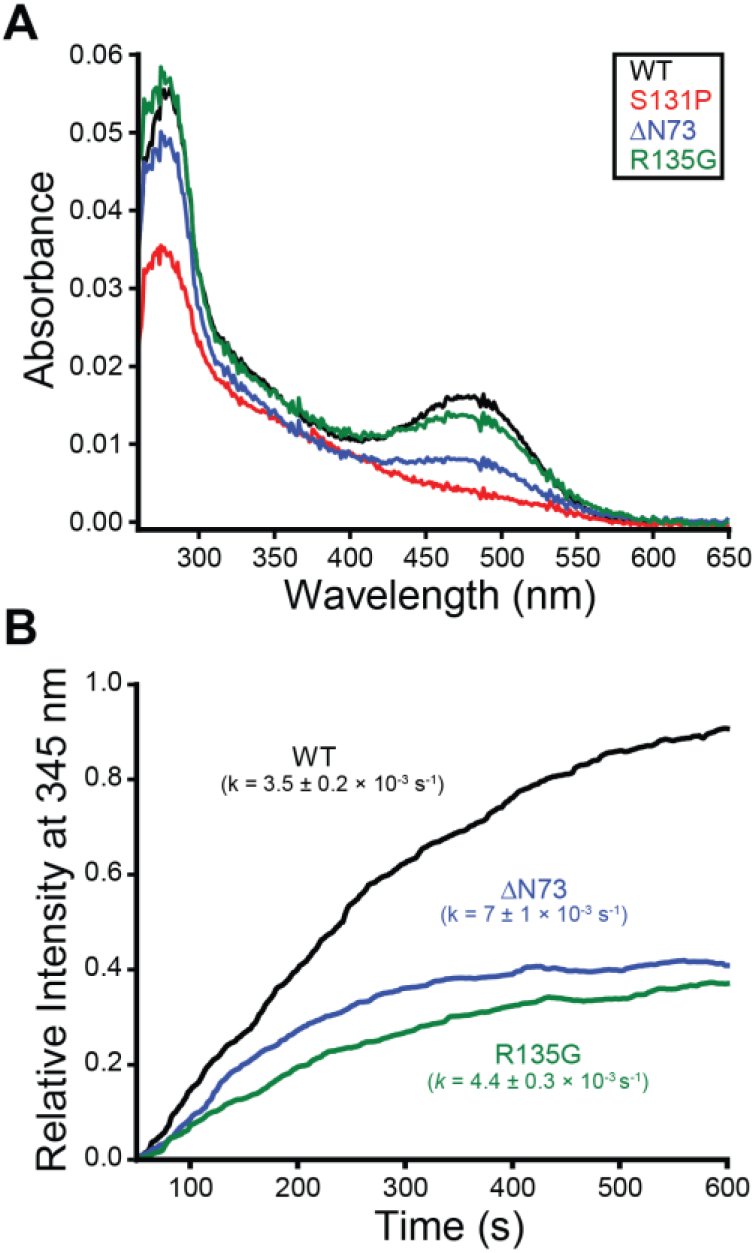
Functional properties of purified retinitis pigmentosa variants. The propensity of purified opsin variants to bind retinal and photoactivate G_t_ are shown. A) Mutant rhodopsin pigments were regenerated in HEK293T cells through the addition of 9-*cis*-retinal then purified into DDM micelles. The relative absorbance of R135G (green), ΔN73 (blue), S131P (red), and WT (black) rhodopsins are plotted as a function of wavelength. B) Purified G_t_ was mixed with regenerated rhodopsins, and the kinetics of G_t_ activation was monitored over time according to the change in fluorescence that arises from nucleotide exchange. The change in fluorescence emission at 345 nm following photoactivation of R135G (green), ΔN73 (blue), and WT (black) rhodopsins are plotted over time. Fitted rate constants are shown for reference.

## Discussion

Corrector molecules hold great promise for the treatment of protein misfolding diseases.(*24*) However, their efficacy varies widely across the spectrum of clinical mutations that enhance misfolding.(*16, 25–28*) Though an understanding of how different variants respond to correctors can aid in their development,(*5*) the sheer volume of clinical variants and the constraints of traditional biochemical and biophysical(*29*) assays has historically precluded their comprehensive characterization. In this work, we utilize DMS to characterize the proteostatic effects of 123 retinopathy-linked rhodopsin variants in HEK293T cells and measure their response to the investigational corrector 9-*cis*-retinal. Though the effects of these mutations may differ in the native context of the rod outer segment, we show that the observed PME of previously characterized variants are highly consistent with previous classifications (Figure 2A) and with the results of another recent high throughput investigation of retinopathy variant expression.(*9*) Our measurements also identify 13 previously uncharacterized variants with severely deficient PME and seven that exhibit moderately deficient PME (Table S1). The 22 other uncharacterized variants characterized herein exhibit robust expression and presumably compromise other aspects of signaling. Our measurements in the presence of retinal reveal that, while variants with the lowest expression exhibit the biggest change in PME (Figure 3 A & C), variants with intermediate expression exhibit the largest absolute increases in PME (Figure 3C). To rationalize these observed mutagenic trends, we outline a thermodynamic framework to interpret the effects of retinal on the PME and to infer which mutants compromise retinal binding. Together, our results provide a holistic overview of the proteostatic effects of known retinopathy variants and identify a discrete subset that are potentially amenable to pharmacological correction.

Our data suggest that 56% of retinopathy variants (69/123) exhibit a considerable attenuation of PME (Table S1)-a figure that is slightly lower than the relative proportion of class II variants associated with other diseases of protein misfolding.(*30–32*) Of these variants, 64% (44/69) exhibit severely reduced PMEs. Consistent with expectations, variants that are predicted to disrupt translocon-mediated membrane integration and/ or to destabilize the native conformation exhibit the largest decreases in expression (Figure 2A). Severe class II variants also exhibit minimal gain in expression in the presence of 5 μM 9-*cis*-retinal (Table S1, Figure 3B). Our thermodynamic projections imply that these mutations are insensitive to retinal because their energetic effects on stability far outweigh the stabilization that is imparted from binding (Figure 4A, Figure S1). Nevertheless, we should note that some of these severe class II variants may also compromise binding. Identification of such variants is complicated by the fact that the magnitude of the expected change in PME is within the error of the experiment (Figure 4). Regardless of the mechanism, variants that fail to respond to correctors are often considered to be “irreversibly” misfolded. However, our thermodynamic framework suggests variants that retain binding affinity should generally exhibit larger gains in PME in the presence of compounds that bind with higher affinity (Figure S1). There is room for optimism in this regard, as 9-*cis*-retinal binds with much lower affinity than rhodopsin’s native cofactor 11-*cis*-retinal (*K*_d_ = 25 pM).(*33*) As is true for other classes of correctors,(*34*) binding affinity is a key consideration for emerging retinoid and non-retinoid rhodopsin correctors. (*10, 14, 35*) Nevertheless, because the observed shifts are rooted in stability, the moderate class II variants that exhibit large shifts in response to 9-*cis*-retinal should generally represent the most favorable target variants for other correctors as well.

Despite many efforts to develop correctors for misfolded rhodopsin variants, there are currently no treatments for adRP or CSNB. Most efforts to discover correctors have been evaluated in relation to their effects on the expression and/ or maturation of the P23H variant-the most common pathogenic rhodopsin variant. (*12–15, 36*) However, P23H is also among the most poorly expressed variants and exhibits only modest sensitivity to retinal (Table S1). Based on this consideration, ongoing corrector screens could potentially achieve better sensitivity by targeting moderate class II variants that exhibit a greater response (i.e. ΔN73, R135G, or Y191C, Table S1). We should also note that the DMS approach described herein could potentially provide an efficient approach to compare the mutation-specific responses to emerging lead compounds. Such experiments will provide key insights as to whether different correctors can rescue distinct variants, or whether there is one common set of “correctable” variants that can be targeted in clinical trials. These considerations highlight new applications of DMS for precision pharmacology.

## Materials and Methods

### Plasmid preparation and mutagenesis

A previously described pcDNA5 vector containing rhodopsin, an N-terminal hemagglutinin (HA) tag, an internal ribosome entry site-dasher GFP cassette, and a Bxb1 recombination site in place of the promoter(*18*) was used to generate a molecular library of barcoded retinopathy variants. We first installed a randomized ten nucleotide “barcode” region upstream of the Bxb1 recombination site using nicking mutagenesis.(*37*) A plasmid preparation containing a mixed population of barcoded vectors was used as a template for 123 individual site-directed mutagenesis reactions to generate a library of retinopathy variants that were found in either the Uniprot Database, the Human Gene Mutation Database, and/ or the Leiden Open Variation Database. Individual clones from each reaction were generated using the GeneJET Plasmid Miniprep Kit (ThermoFisher Scientific, Waltham, MA) and sequenced to confirm the sequence of each mutated open reading frame and to determine its corresponding 10 base barcode sequence. Plasmids encoding individual variants were pooled and electroporated into electro-competent NEB10β cells (New England Biolabs, Ipswitch, MA), which were then grown in liquid culture overnight and purified using the ZymoPure endotoxin-free midiprep kit (Zymo Research, Irvine, CA). The Bxb1 recombinase expression vector (pCAG-NLS-HA Bxb1) was provided by Douglas Fowler.

### Production and fractionation of recombinant cell lines

A pool of recombinant stable cells expressing individual retinopathy variants was generated using a previously described stable HEK293T cell line containing a genomic Tet-Bxb1-BFP landing pad.(*17*) Recombinant cells were generated and isolated as was previously described.(*18*) Briefly, cells grown in 10 cm dishes in complete Dulbecco’s modified Eagle medium (Gibco, Carlsbad, CA) supplemented with 10% fetal bovine serum (Corning, Corning, NY) and penicillin (100 U/ml)/streptomycin (100 μg/ml) (complete media) were co-transfected with our library of retinopathy variants and the Bxb1 recombinase expression vector using Fugene 6 (Promega, Madison, WI). Doxycycline (2 μg/mL) was added one day after transfection and the cells were grown at 33 °C for the following 4 days. On the 4^th^ day, cells were sorted using the BD FACS Aria II (BD Biosciences, Franklin Lakes, NJ) to isolate GFP positive/ BFP negative cells that had undergone recombination. These cells were grown in 10 cm dishes with complete media supplemented with doxycycline (2 μg/mL) for up to seven days. Where indicated, cells were incubated with 5 μM 9-*cis*-retinal for 16 hours prior to sorting. Rhodopsin expressed at the plasma membrane of recombinant cells was labeled with a DyLight 550–conjugated anti-HA antibody (ThermoFisher, Waltham, MA). Labelled cells were then fractionated into quartiles according to surface immunostaining intensity using a FACS Aria IIu fluorescence activated cell sorter (BD Biosciences, Franklin Lakes, NJ). At least 2 million cells from each fraction were isolated to ensure exhaustive sampling. Fractionated sub-populations were expanded in 10 cm culture dishes prior to harvesting and freezing 10-20 million cells per quartile fraction for the downstream genetic analysis.

### Extraction of genomic DNA and preparation of next-generation sequencing libraries

To track the expression of individual retinopathy variants, we first extracted the genomic DNA (gDNA) from each cellular fraction using the GenElute Mammalian Genomic DNA Miniprep kit (Sigma-Aldrich, St. Louis, MO). A previously described semi-nested polymerase chain reaction (PCR) technique(*17*) was then used to selectively amplify the barcoded region of the recombined plasmids within the gDNA. Briefly, an initial PCR reaction was used to first amplify the region of interest from the gDNA. The product of this reaction was then used as a template for a second PCR reaction that amplified the barcoded region while installing indexed Illumina adapter sequences. Amplicons were gel-purified using the Zymoclean Gel DNA Recovery Kit (Zymo Research, Irvine, CA). The purity of each sequencing library was confirmed using an Agilent 2200 TapeStation (Agilent Technologies, Santa Clara, CA). Libraries were sequenced using a NextSeq 500 Mid Output 150-cycle kit (Illumina, San Diego, CA) at an average depth of ~2 million reads per quartile.

### Estimation of surface immunostaining levels from deep mutational scanning data

Surface immunostaining levels were estimated from sequencing data using a computational approach described previously.(*18*) Briefly, low quality reads that were likely to contain more than one error were removed from the analysis. The remaining reads containing one of the 123 barcodes corresponding to a variant of interest were then rarefied to generate subsampled datasets with a uniform number of reads for each sample. We then calculated weighted-average immunostaining intensity values for each barcode/ variant using the following equation:

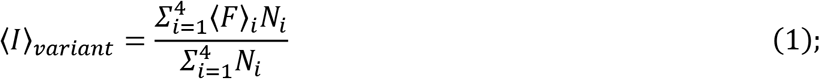

where 〈I〉_variant_ is the weighted-average fluorescence intensity of a given variant, 〈F〉_i_ is the mean fluorescence intensity associated with cells from the i_th_ FACS quartile, and Ni is the number of barcode/ variant reads in the i_th_ FACS quartile. Variant intensities from each replicate were normalized relative to one another using the mean surface immunostaining intensity of the recombinant cell population on each day to account for small variations in laser power and/ or detector voltage. Intensity values reported herein represent the average normalized intensity values from three replicate experiments.

### Derivation of thermodynamic bounds for variant expression

Estimates for the upper and lower bounds for the change in variant expression in the presence of 9-*cis*-retinal were derived based on a series of simplifying assumptions concerning the relationship between the folding energetics, binding energetics, and the expression of the mature protein at the plasma membrane. The plasma membrane expression of integral membrane proteins in eukaryotic cells should generally scale with their thermodynamic preference for their native fold.(*21, 31, 38*) Therefore, we first assume that surface immunostaining is proportional to the combined equilibrium fraction of folded opsin and rhodopsin (*f*_fold_). We also assume that this collection of variants includes some that are stable (ΔG_fold_ ≤ −3 kcal/ mol) and others that are unstable (ΔG_fold_ ≥ 3 kcal/ mol). Based on this criterion, the highest and lowest variant immunostaining intensities were taken as the signal generated by cells expressing fully folded (*f*_fold_ ~ 1) and fully unfolded (*f*_fold_ ~ 0) variants, respectively. This scaling can be used to approximate the fraction of folded protein (*f*_fold_) using the following generalizable equation:

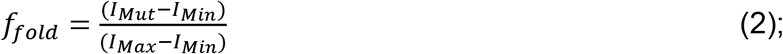

where *I*_Mut_ is the observed immunostaining intensity of the variant of interest, *I*_Max_ is the highest observed variant immunostaining intensity, and *I*_Min_ is the lowest observed variant immunostaining intensity under a given condition. The approximated fraction of folded apoprotein (*f*_fold,Apo_) can be calculated for each variant by plugging observed intensity values in the absence of retinal (*I*_Mut,Apo_, *I*_Max,Apo_, and *I*_Min,Apo_) into Equation 2. Likewise, the approximated fraction of folded protein in the presence of retinal (*f*_fold,Ret_) can be calculated for each variant by plugging observed intensity values in the presence of retinal (*I*_Mut,Ret_, *I*_Max,Ret_, and *I*_Min,Ret_) into Equation 2.

For mutations that fully compromise binding (*K*_d,Mut_ → ∞), *f*_fold_ should be the same in the presence and absence of retinal. Setting *f*_fold,Ret_ equal to *f*_fold,Apo_ and solving for *I*_Mut,Ret_ produces the following equation relating the projected *I*_Mut,Ret_ for variants that fail to bind retinal to the corresponding *I*_Mut,Apo_ and the intensity limits within each experiment:

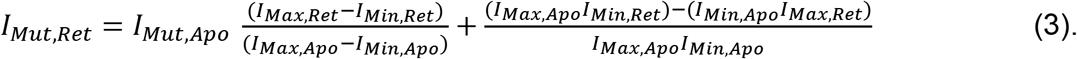

Equation 3 can be used to derive a lower boundary for the change in immunostaining intensity for arbitrary variants in the presence of retinal based on the differences in the observed experimental fluorescence intensities (red dashes, Figure 4A).

Approximations of *f*_fold_ must be cast in terms of the free energy of folding (ΔG_fold_) to project the effects of binding energetics on the immunostaining intensities of variants that bind retinal. Approximated *f*_fold_ values can then used to calculate the corresponding value of ΔG_fold_ using the following generalizable equation:

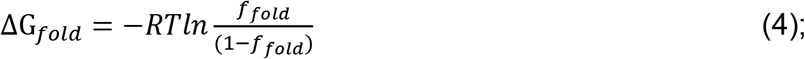

where R is the gas constant, and T is the temperature. The free energy of folding in the absence of retinal (ΔG_Apo_) for each variant can be determined by plugging *f*_fold,Apo_ into Equation 4. The ΔG_Apo_ value for each variant can then be used to estimate the apparent free energy of folding in the presence of retinal (ΔG_App_) using the following previously derived equation:(*39*)

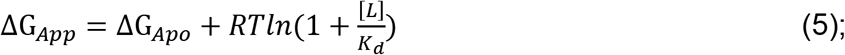

where [L] is the concentration of the retinal ligand, and *K*_d_ is the equilibrium dissociation constant for retinal. The second term of Equation 5 can represents the expected change in the free energy of folding in the presence of retinal (ΔΔG_fold_), which works out to −1.1 kcal/ mol for variants that do not perturb 9-*cis*-retinal binding (*K*_d_ ~ 0.9 μM)(*14*) in the presence of 5 μM 9-*cis*-retinal. Using Equation 4 to re-cast the ΔG_Apo_ and ΔG_App_ terms in Equation 5 in terms of *f*_fold,Apo_ and *f*_fold,App_ results in the following equation describing the extent to which retinal should increase the fraction of folded protein:

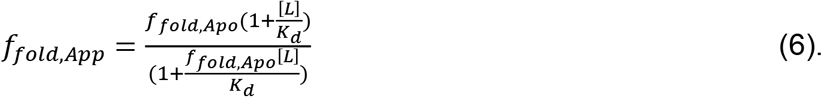

Together, these equations can be used to project how *I*_Mut,Ret_ should depend on *I*_Mut,Apo_, [L], and *K*_d_ as follows. Plugging observed intensity values in the presence of retinal (*I*_Mut,Ret_, *I*_Max,Ret_, and *I*_Min,Ret_) into Equation 2 and solving for *I*_Mut,Ret_ yields the following equation:

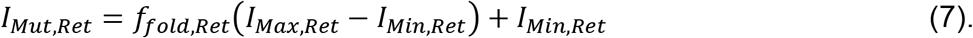

Combining Equations 6 and 7 then results in the following:

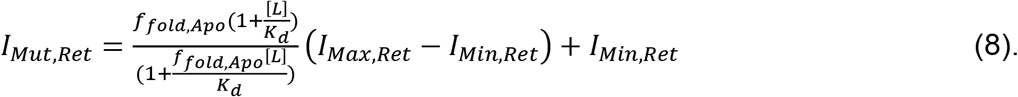

Finally, Equation 2 can be used to re-cast *f*_fold,Apo_ in terms of immunostaining intensities as follows:

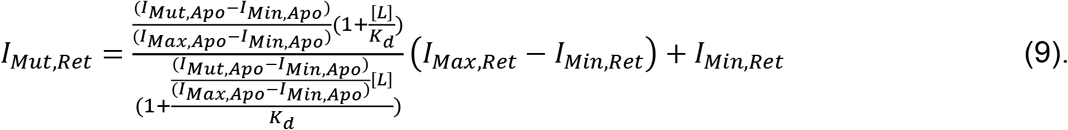

Equation 9 can be used to derive an upper bound for the *I*_Mut,Ret_ of variants that do not perturb binding (*K*_d,Mut_ = *K*_d,WT_ = 0.9 μM) in the presence of the experimental dosage of retinal (5 μM, blue dashes, Figure 4A).

Most variants fall between these upper and lower bounds (Figure 4A). If it is assumed that the difference between this projected *I*_Mut,Ret_ and the observed *I*_Mut,Ret_ for a given variant arises solely from the effects of the mutation on binding energetics, then an approximation for *K*_d,Mut_ can be projected from observed immunostaining intensities as follows. First, Equation 2 can be used to calculate *f*_fold,Ret_ from, *I*_Max,Ret_, *I*_Min,Ret_, and the observed Immunostaining intensity of the variant in the presence of retinal. The corresponding ΔG_app_ can then be determined by plugging *f*_fold,Ret_ into Equation 3. An approximated *K*_d,Mut_ can then be calculated by plugging the resulting ΔG_app_ along with the ΔG_fold_ for the apo form of the variant and [L] into Equation 5, then solving for *K*_d_.

### Computational predictions of the effects of mutations on folding energetics

The impacts of mutations on the thermodynamic stability of rhodopsin were estimated by constructing molecular models of each rhodopsin variant and comparing their stability using a membrane protein-specific Rosetta energy function as described previously.(*18*) Briefly, a high-resolution crystal structure of bovine rhodopsin [Protein Data Bank (PDB) 3C9L] was used to generate homology models of each human rhodopsin variant as was previously described.(*21*) A Rosetta ΔΔG protocol was then using to estimate the effects of each mutant on the conformational stability of the native fold.(*40*) The effect of RP mutations on cotranslational TM domain integration was estimated using a previously described ΔG predictor algorithm (https://dgpred.cbr.su.se/).(*41*)

### Computational predictions of the effects of mutations on binding energetics

We estimated the effects of mutations on the binding energy using a series of structural models for each mutant in both its apo state (opsin) and its covalently-bound state (rhodopsin). Opsin and rhodopsin structures were modeled based on a crystal structure of bovine rhodopsin (PDB 2PED), which features a linkage to the 9-*cis*-retinal isomer used in these studies. To facilitate Rosetta modeling, the covalently linked retinal was treated as a non-canonical amino acid (NCAA) by converting its SDF to a molfile with Open Babel. The Biochemical Library (BCL) code repository (http://www.meilerlab.org/index.php/bcl_commons/show/b_apps_id/1) was then used to generate conformer libraries and Rosetta params files. The MutateResidue mover was then used to mutate K296 to this NCAA representation to facilitate its correct recognition by Rosetta. We used Rosetta scripts to introduce each mutation into the WT model, then used FastRelax in dualspace with the ref2015_cart energy function to refine each model. A custom packer palette was included to expand the default type sets used during relaxation of the retinal NCAA. To estimate the effects of mutations on the initial (non-covalent) binding reaction, we first converted the retinal-conjugated K296 in the PDB 2PED model back to lysine. We then used RosettaLigand to dock a non-covalent 9-*cis*-retinal back into the pocket and used the resulting structure as a template to generate variant models using the cartesian_ddg interface in Rosetta. The resulting models were then relaxed without constraints and the free energy of binding for the lowest energy structure was then calculated using the K_DEEP_ web server.(*23*) To estimate the effects of mutations on the free energy of the covalently-bound structure, the interface energy between retinal and the portions of the protein was estimated by taking the average of the InteractionEnergyMetric of the 5 lowest energy models for each mutant rhodopsin structure.

### Purification and Spectroscopic Characterization of Rhodopsin Variants

Rhodopsin pigments were purified using a modified version of a previously described protocol.(*42*) Briefly, polyethyleneimine was used to transfect twenty 10 cm dishes of HEK293T cells with pcDNA5 expression vectors containing the cDNA for human WT, S131P, R135G, and ΔN73 rhodopsin. Cells were treated overnight with 7.5 μM 9-*cis*-retinal beginning 24 hours post-transfection, and were grown in the dark to facilitate the regeneration of each pigment. Cells were harvested 48 hours post-transfection under dim red light, then pelleted by centrifugation at 800 g. Cell pellets were either stored at −80 °C or were directly lysed by rotating the slurry in the dark for 1 h at 4 °C in 20 mM Bis-tris propane (BTP, pH 7.5) containing 120 mM NaCl, 20 mM n-dodecyl-β-D-maltopyranoside (DDM), and a protease inhibitor cocktail. Lysates were then clarified by centrifugation at 100,000 g for one hour at 4 °C. Pigments were purified from the supernatant using a 1D4 anti-rhodopsin immuno-affinity chromatography. 200 μL of 1D4 resin (6 mg 1D4 monoclonal anti-rhodopsin antibody/ml agarose beads) were added to the supernatant and rotated for one hour at 4 °C. The beads were then transferred to a column and washed with 12 mL of 20 mM BTP (pH 7.5) containing 120 mM NaCl and 2 mM DDM, followed by a wash with 20 mM BTP (pH 7.5) containing 500 mM NaCl and 2 mM DDM. Rhodopsin pigments were eluted with 20 mM BTP (pH 7.5) containing 120 mM NaCl, 2 mM DDM, and 0.6 mg/ml of an elution peptide (TETSQVAPA).(*21, 43, 44*) The UV-visible spectra of purified rhodopsin pigments were recorded in the dark using a Cary 60 UV-visible spectrophotometer (Varian, Palo Alto, CA). The concentrations of purified pigments regenerated with 9-*cis*-retinal rhodopsins were determined assuming a molar extinction coefficient of ε_485nm_ = 43,600 M^−1^cm^−1^.(*45*)

### G_t_ activation measurements

The ability of mutant pigments to activate G_t_ *in vitro* was measured as previously described.(*21*) Briefly, G_t_ was extracted and purified from frozen rod outer segment membranes isolated from 100 dark-adapted bovine retinas.(*15, 44*) Purified G_t_ was mixed with purified rhodopsin variants to final concentrations of 250 nM and 25 nM, respectively, in 20 mM BTP (pH 7.0) containing 120 mM NaCl, 1 mM MgCl_2_ and 1 mM DDM. This mixture was then illuminated for 1 min with a Fiber-Light illuminator (Dolan Jenner Industries Inc., Boxborough, MA) through a 480-520 nm band-pass wavelength filter (Chroma Technology Corporation, Bellows Falls, VT) in order to photoactivate the rhodopsin pigments. 10 μM GTPγS was then added following illumination, and the change in tryptophan fluorescence associated with the exchange of guanyl nucleotides within the α subunit of G_t_ was measured for 1200 s with a FL 6500 Fluorescence Spectrometer (PerkinElmer, Waltham, MA). Excitation and emission wavelengths were set at 300 nm and 345 nm, respectively.(*15, 43*) G_t_ activation rates were determined by fitting the change in fluorescence intensity over the initial 600 s with a single exponential function.

## Acknowledgements

We thank Christiane Hassel and the Indiana University Flow Cytometry Core Facility for technical support. We thank the Indiana University Center for Genomics and Bioinformatics for experimental support. This research was supported by grants from the National Institutes of Health (NIH) (R01GM129261 to J. P. S., R01EY025214 to B. J., R01HL122010 to J. M., R01GM080403 to J. M., and R01DA046138 to J. M.). F. J. R. acknowledges receipt of a predoctoral fellowship from the Graduate Training Program in Quantitative and Chemical Biology at Indiana University (T32 GM109825). The authors also thank the Visual Science Research Center Core at Case Western Reserve University (supported by NIH grant P30EY011373).

## References

1. C. Sanders, J. Myers, Disease-related misassembly of membrane proteins. Annu Rev Biophys Biomol Struct 33, 25–51 (2004).

2. J. T. Marinko et al., Folding and Misfolding of Human Membrane Proteins in Health and Disease: From Single Molecules to Cellular Proteostasis. Chem Rev 119, 5537–5606 (2019).

3. J. P. Schlebach, C. R. Sanders, The safety dance: biophysics of membrane protein folding and misfolding in a cellular context. Q Rev Biophys 48, 1–34 (2015).

4. B. M. Kroncke, C. G. Vanoye, J. Meiler, A. L. George, Jr., C. R. Sanders, Personalized biochemistry and biophysics. Biochemistry 54, 2551–2559 (2015).

5. G. Veit et al., From CFTR biology toward combinatorial pharmacotherapy: expanded classification of cystic fibrosis mutations. Mol Biol Cell 27, 424–433 (2016).

6. P. G. Middleton et al., Elexacaftor-Tezacaftor-Ivacaftor for Cystic Fibrosis with a Single Phe508del Allele. New Eng J Med 381, 1809–1819 (2019).

7. D. Athanasiou et al., The molecular and cellular basis of rhodopsin retinitis pigmentosa reveals potential strategies for therapy. Prog Retin Eye Res 62, 1–23 (2018).

8. S. Kaushal, H. G. Khorana, Structure and function in rhodopsin. 7. Point mutations associated with autosomal dominant retinitis pigmentosa. Biochemistry 33, 6121–6128 (1994).

9. A. Wan, E. Place, E. A. Pierce, J. Comander, Characterizing variants of unknown significance in rhodopsin: A functional genomics approach. Human Mutation 40, 1127–1144 (2019).

10. P. Behnen et al., A Small Chaperone Improves Folding and Routing of Rhodopsin Mutants Linked to Inherited Blindness. iScience 4, 1–19 (2018).

11. C. Zeitz et al., Identification and functional characterization of a novel rhodopsin mutation associated with autosomal dominant CSNB. Invest Ophthalmol Vis Sci 49, 4105–4114 (2008).

12. S. M. Noorwez et al., Retinoids assist the cellular folding of the autosomal dominant retinitis pigmentosa opsin mutant P23H. J Biol Chem 279, 16278–16284 (2004).

13. K. Ohgane, K. Dodo, Y. Hashimoto, Retinobenzaldehydes as proper-trafficking inducers of folding-defective P23H rhodopsin mutant responsible for retinitis pigmentosa. Bioorg Med Chem 18, 7022–7028 (2010).

14. Y. Chen et al., A novel small molecule chaperone of rod opsin and its potential therapy for retinal degeneration. Nat Commun 9, 1976 (2018).

15. J. T. Ortega, T. Parmar, B. Jastrzebska, Flavonoids enhance rod opsin stability, folding, and self-association by directly binding to ligand-free opsin and modulating its conformation. J Biol Chem 294, 8101–8122 (2019).

16. M. P. Krebs et al., Molecular mechanisms of rhodopsin retinitis pigmentosa and the efficacy of pharmacological rescue. J Mol Biol 395, 1063–1078 (2010).

17. K. A. Matreyek, J. J. Stephany, D. M. Fowler, A platform for functional assessment of large variant libraries in mammalian cells. Nucleic Acids Res 45, e102 (2017).

18. W. D. Penn et al., Probing biophysical sequence constraints within the transmembrane domains of rhodopsin by deep mutational scanning. Sci Adv 6, eaay7505 (2020).

19. L. Zhang et al., Contribution of hydrophobic and electrostatic interactions to the membrane integration of the Shaker K+ channel voltage sensor domain. Proc Natl Acad Sci U S A 104, 8263–8268 (2007).

20. A. G. McKee et al., Systematic profiling of temperature- and retinal-sensitive rhodopsin variants by deep mutational scanning. J Biol Chem 297, 101359 (2021).

21. F. J. Roushar et al., Contribution of Cotranslational Folding Defects to Membrane Protein Homeostasis. J Am Chem Soc 141, 204–215 (2019).

22. S. A. Combs et al., Small-molecule ligand docking into comparative models with Rosetta. Nat Protoc 8, 1277–1298 (2013).

23. J. Jimenez, M. Skalic, G. Martinez-Rosell, G. De Fabritiis, KDEEP: Protein-Ligand Absolute Binding Affinity Prediction via 3D-Convolutional Neural Networks. J Chem Inf Model 58, 287–296 (2018).

24. F. Chiti, J. W. Kelly, Small molecule protein binding to correct cellular folding or stabilize the native state against misfolding and aggregation. Curr Opin Struct Biol 72, 267–278 (2022).

25. G. Veit et al., Allosteric folding correction of F508del and rare CFTR mutants by elexacaftor-tezacaftor-ivacaftor (Trikafta) combination. JCI Insight 5, (2020).

26. S. T. Han et al., Residual function of cystic fibrosis mutants predicts response to small molecule CFTR modulators. JCI Insight 3, (2018).

27. G. Veit et al., A Precision Medicine Approach to Optimize Modulator Therapy for Rare CFTR Folding Mutants. J Pers Med 11, (2021).

28. G. Veit et al., Structure-guided combination therapy to potently improve the function of mutant CFTRs. Nat Med 24, 1732–1742 (2018).

29. K. G. Fleming, Energetics of membrane protein folding. Annu Rev Biophys 43, 233–255 (2014).

30. Z. Wang, J. Moult, SNPs, protein structure, and disease. Human mutation 17, 263–270 (2001).

31. J. P. Schlebach et al., Conformational Stability and Pathogenic Misfolding of the Integral Membrane Protein PMP22. J Am Chem Soc 137, 8758–8768 (2015).

32. H. Huang et al., Mechanisms of KCNQ1 channel dysfunction in long QT syndrome involving voltage sensor domain mutations. Sci Adv 4, eaar2631 (2018).

33. H. Tian, T. P. Sakmar, T. Huber, The Energetics of Chromophore Binding in the Visual Photoreceptor Rhodopsin. Biophys J 113, 60–72 (2017).

34. J. A. Janovick et al., Structure-activity relations of successful pharmacologic chaperones for rescue of naturally occurring and manufactured mutants of the gonadotropin-releasing hormone receptor. J Pharmacol Exp Ther 305, 608–614 (2003).

35. D. Mattle et al., Ligand channel in pharmacologically stabilized rhodopsin. Proc Natl Acad Sci U S A 115, 3640–3645 (2018).

36. J. T. Ortega, T. Parmar, M. Carmena-Bargueno, H. Perez-Sanchez, B. Jastrzebska, Flavonoids improve the stability and function of P23H rhodopsin slowing down the progression of retinitis pigmentosa in mice. J Neurosci Res, (2022).

37. E. E. Wrenbeck et al., Plasmid-based one-pot saturation mutagenesis. Nat Methods 13, 928–930 (2016).

38. R. L. Wiseman, E. T. Powers, J. N. Buxbaum, J. W. Kelly, W. E. Balch, An adaptable standard for protein export from the endoplasmic reticulum. Cell 131, 809–821 (2007).

39. C. Park, S. Marqusee, Pulse proteolysis: a simple method for quantitative determination of protein stability and ligand binding. Nat Methods 2, 207–212 (2005).

40. E. H. Kellogg, A. Leaver-Fay, D. Baker, Role of conformational sampling in computing mutation-induced changes in protein structure and stability. Proteins 79, 830–838 (2011).

41. T. Hessa et al., Molecular code for transmembrane-helix recognition by the Sec61 translocon. Nature 450, 1026–1030 (2007).

42. Y. Chen, H. Tang, High-throughput screening assays to identify small molecules preventing photoreceptor degeneration caused by the rhodopsin P23H mutation. Methods Mol Biol 1271, 369–390 (2015).

43. D. P. Mallory et al., The Retinitis Pigmentosa-Linked Mutations in Transmembrane Helix 5 of Rhodopsin Disrupt Cellular Trafficking Regardless of Oligomerization State. Biochemistry 57, 5188–5201 (2018).

44. B. Jastrzebska, Oligomeric state of rhodopsin within rhodopsin-transducin complex probed with succinylated concanavalin A. Methods Mol Biol 1271, 221–233 (2015).

45. J. D. Spalink, A. H. Reynolds, P. M. Rentzepis, W. Sperling, M. L. Applebury, Bathorhodopsin intermediates from 11-cis-rhodopsin and 9-cis-rhodopsin. Proc Natl Acad Sci U S A 80, 1887–1891 (1983).

