## Supplemental Material for "Molecular Basis for Variations in the Sensitivity of Pathogenic Rhodopsin Variants to 9-*cis*-Retinal"

**Contents:**
-Figure S1
-Table S1

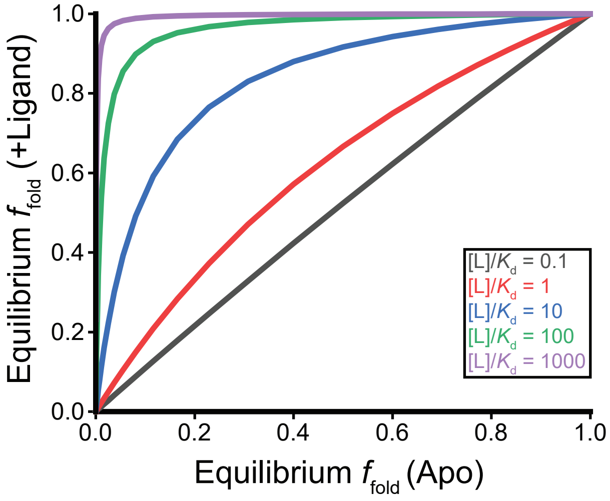


**Figure S1. Thermodynamic Relationship Between Ligand Binding on the Equilibrium Fraction of Folded Protein.** Equation 5 was used to similate the impact of ligand binding on the equilibrium fraction of folded protein as a function of the ratio of ligand to its equilibrium dissociation constant. The equilibrium fraction of folded protein in the presence of ligand is plotted as a function of corresponding equilibrium fraction of folded apoprotein. This simulation reveals how the degree of rescue offered by a corrector should relate to the thermodynamic stability of the variant (x-coordinate), the concentration of the ligand, and its binding affinity.

**Table S1. Classification, Retinal-Sensitivity, and Plasma Membrane Expression (PME) of Mutations in Rhodopsin**

| Mutation | Prior Classification | DMS Classification | Surface Immunostaing Intensity (Apo)* | Surface Immunostaing Intensity (+Ret)* |
|---|---|---|---|---|
| T4K | Class 2[1, 2] | Non-Class 2 | 21459 ± 1373 | 19838 ± 3633 |
| P12R | Unclassified | Moderate Class 2 | 3044 ± 30 | 5245 ± 834 |
| N15S | Class 2[1] | Severe Class 2 | 2058 ± 88 | 2473 ± 303 |
| T17M | Class 2[1-3] | Severe Class 2 | 2211 ± 75 | 2604 ± 227 |
| V20G | Class 2[1] | Severe Class 2 | 2251 ± 91 | 2774 ± 268 |
| P23H | Class 2[1-3] | Severe Class 2 | 1716 ± 7 | 2133 ± 171 |
| P23L | Class 2[1] | Severe Class 2 | 1585 ± 55 | 1988 ± 255 |
| P23A | Class 2[1] | Severe Class 2 | 2295 ± 24 | 2920 ± 275 |
| Q28H | Class 2[1, 2] | Severe Class 2 | 1905 ± 82 | 2236 ± 102 |
| Q28R | Class 2[1] | Severe Class 2 | 1826 ± 51 | 2203 ± 117 |
| M39R | Class 4[1] | Moderate Class 2 | 7406 ± 61 | 8483 ± 646 |
| M44T | Class 5[1] | Non-Class 2 | 20520 ± 250 | 22775 ± 1128 |
| F45L | Class 7[1] | Non-Class 2 | 21466 ± 261 | 23900 ± 1331 |
| L46R | Unclassified | Severe Class 2 | 1575 ± 55 | 1887 ± 86 |
| L47R | Class 2[4] | Moderate Class 2 | 2891 ± 92 | 5580 ± 159 |
| G51R | Class 2[1, 2, 4] | Moderate Class 2 | 6840 ± 236 | 11251 ± 1695 |
| G51A | Unclassified | Non-Class 2 | 19022 ± 278 | 21907 ± 945 |
| G51V | Class 2[1, 2] | Non-Class 2 | 19866 ± 375 | 21197 ± 624 |
| F52Y | Unclassified | Moderate Class 2 | 12693 ± 203 | 16135 ± 2076 |
| P53R | Class 2[1, 2] | Severe Class 2 | 1659 ± 75 | 3510 ± 247 |
| F56Y | Unclassified | Non-Class 2 | 16591 ± 109 | 18485 ± 1691 |
| L57R | Class 2[4] | Moderate Class 2 | 2956 ± 128 | 4650 ± 184 |
| T58R | Class 2[1, 3, 4] | Non-Class 2 | 17889 ± 209 | 22138 ± 1739 |
| R69H | Unclassified | Non-Class 2 | 20184 ± 589 | 23143 ± 999 |
| del N73 | Unclassified | Moderate Class 2 | 4913 ± 293 | 11881 ± 2355 |
| N78I | Class 2[4] | Moderate Class 2 | 3185 ± 274 | 4998 ± 625 |
| V87L | Unclassified | Non-Class 2 | 18186 ± 779 | 22577 ± 1112 |
| V87D | Class 2[1-3] | Moderate Class 2 | 3731 ± 472 | 5216 ± 642 |
| L88P | Class 2[4] | Moderate Class 2 | 7799 ± 124 | 12400 ± 2582 |
| G89D | Class 2[1-4] | Moderate Class 2 | 10580 ± 514 | 9423 ± 1590 |
| G90D | Class 6[1] | Non-Class 2 | 21297 ± 242 | 20572 ± 503 |
| G90V | Class 6[1] | Non-Class 2 | 21101 ± 268 | 20872 ± 1098 |
| T94I | Class 6[1] | Non-Class 2 | 25614 ± 185 | 24016 ± 497 |
| V104I | Unclassified | Non-Class 2 | 20746 ± 333 | 23099 ± 1134 |
| V104F | Unclassified | Non-Class 2 | 17108 ± 351 | 18678 ± 550 |
| G106R | Class 2[1, 2] | Moderate Class 2 | 6597 ± 210 | 8280 ± 1114 |
| G106W | Class 2[1-3] | Severe Class 2 | 2101 ± 99 | 2993 ± 315 |
| G109R | Class 2[4] | Moderate Class 2 | 14723 ± 150 | 13588 ± 2744 |
| C110R | Class 2[1] | Severe Class 2 | 1645 ± 73 | 2062 ± 114 |
| C110Y | Class 2[1, 2] | Severe Class 2 | 1660 ± 86 | 2036 ± 96 |
| C110F | Class 2[1] | Severe Class 2 | 1571 ± 80 | 1920 ± 127 |
| G114D | Unclassified | Severe Class 2 | 1663 ± 103 | 1958 ± 81 |
| G114V | Unclassified | Severe Class 2 | 1575 ± 41 | 2544 ± 155 |
| E122G | Unclassified | Non-Class 2 | 21147 ± 314 | 22740 ± 1211 |

| Mutation | Prior Classification | DMS Classification | Surface Immunostaing Intensity (Apo)* | Surface Immunostaing Intensity (+Ret)* |
|---|---|---|---|---|
| L125R | Class 2[1] | Non-Class 2 | 19889 ± 220 | 18451 ± 1729 |
| S127F | Unclassified | Moderate Class 2 | 2693 ± 178 | 3591 ± 326 |
| L131P | Unclassified | Severe Class 2 | 1882 ± 130 | 3211 ± 325 |
| R135P | Class 3[1] | Moderate Class 2 | 8559 ± 202 | 13039 ± 2177 |
| R135L | Class 2[3] | Moderate Class 2 | 4085 ± 130 | 7931 ± 378 |
| R135G | Class 3[1] | Moderate Class 2 | 8791 ± 181 | 14820 ± 2169 |
| R135W | Class 2[3] | Severe Class 2 | 2129 ± 65 | 7701 ± 299 |
| V137M | Class 5[1] | Non-Class 2 | 20724 ± 216 | 22958 ± 621 |
| C140S | Unclassified | Non-Class 2 | 20898 ± 309 | 23562 ± 1372 |
| E150K | Unclassified | Non-Class 2 | 20855 ± 324 | 23140 ± 993 |
| T160T | Unclassified | Non-Class 2 | 20350 ± 359 | 22847 ± 1185 |
| W161R | Class 2[1] | Severe Class 2 | 1812 ± 54 | 2330 ± 231 |
| A164E | Class 2[1] | Severe Class 2 | 1603 ± 106 | 2606 ± 219 |
| A164V | Class 2[1] | Non-Class 2 | 18661 ± 308 | 20327 ± 713 |
| C167R | Class 2[1, 2, 4] | Non-Class 2 | 20651 ± 320 | 23214 ± 1035 |
| C167W | Class 2[1, 4] | Severe Class 2 | 1904 ± 28 | 3093 ± 197 |
| A169P | Class 2[4] | Severe Class 2 | 1475 ± 67 | 2386 ± 119 |
| P170R | Unclassified | Severe Class 2 | 1461 ± 10 | 1926 ± 283 |
| P171E | Unclassified | Severe Class 2 | 1738 ± 91 | 2465 ± 117 |
| P171Q | Class 2[1] | Severe Class 2 | 1675 ± 74 | 2429 ± 123 |
| P171L | Class 2[1, 2] | Severe Class 2 | 1874 ± 87 | 2679 ± 177 |
| G174S | Unclassified | Non-Class 2 | 19427 ± 265 | 23140 ± 82 |
| S176F | Unclassified | Severe Class 2 | 1649 ± 36 | 2045 ± 212 |
| Y178N | Class 2[1] | Severe Class 2 | 1622 ± 73 | 2916 ± 178 |
| Y178D | Class 2[1] | Severe Class 2 | 1604 ± 83 | 2142 ± 38 |
| I179F | Class 2[4] | Moderate Class 2 | 4667 ± 55 | 7646 ± 964 |
| P180A | Class 2[4] | Moderate Class 2 | 3883 ± 159 | 6555 ± 480 |
| P180S | Class 2[4] | Severe Class 2 | 1667 ± 76 | 2127 ± 123 |
| E181K | Class 2[1, 2] | Severe Class 2 | 1705 ± 42 | 2236 ± 25 |
| G182S | Class 2[1] | Moderate Class 2 | 3154 ± 213 | 5613 ± 471 |
| Q184P | Unclassified | Severe Class 2 | 1852 ± 41 | 2236 ± 126 |
| C185R | Class 2[1] | Severe Class 2 | 1816 ± 28 | 2271 ± 319 |
| S186P | Class 2[2] | Severe Class 2 | 1582 ± 51 | 1991 ± 137 |
| S186W | Class 2[4] | Non-Class 2 | 27766 ± 317 | 28230 ± 875 |
| C187Y | Class 2[1] | Severe Class 2 | 1876 ± 64 | 2166 ± 54 |
| G188R | Class 2[1, 2] | Severe Class 2 | 1660 ± 66 | 2113 ± 289 |
| D190N | Class 2[1] | Moderate Class 2 | 4182 ± 24 | 5586 ± 797 |
| D190G | Class 2[1, 3] | Severe Class 2 | 1788 ± 76 | 2442 ± 115 |
| D190Y | Class 2[1, 2] | Severe Class 2 | 1757 ± 98 | 2216 ± 130 |
| Y191C | Unclassified | Non-Class 2 | 17171 ± 396 | 22547 ± 458 |
| T193M | Class 2[4] | Moderate Class 2 | 12826 ± 604 | 17646 ± 1170 |
| P196T | Unclassified | Non-Class 2 | 19201 ± 404 | 22146 ± 1183 |
| I205S | Unclassified | Non-Class 2 | 16477 ± 104 | 19163 ± 538 |
| M207K | Unclassified | Non-Class 2 | 18429 ± 352 | 17930 ± 1375 |
| M207R | Class 2[4] | Non-Class 2 | 15790 ± 243 | 13919 ± 804 |

| Mutation | Prior Classification | DMS Classification | Surface Immunostaing Intensity (Apo)* | Surface Immunostaing Intensity (+Ret)* |
|---|---|---|---|---|
| V209M | Class 7[1] | Non-Class 2 | 18653 ± 98 | 21664 ± 690 |
| V210F | Unclassified | Non-Class 2 | 19755 ± 141 | 22284 ± 776 |
| H211R | Class 2[1] | Severe Class 2 | 1842 ± 5 | 2237 ± 181 |
| I214N | Class 2[4] | Non-Class 2 | 17234 ± 484 | 19082 ± 1133 |
| P215T | Unclassified | Severe Class 2 | 1521 ± 78 | 1972 ± 215 |
| P215L | Unclassified | Severe Class 2 | 2307 ± 112 | 3037 ± 207 |
| M216K | Class 2[4] | Non-Class 2 | 17663 ± 243 | 20170 ± 705 |
| M216L | Unclassified | Non-Class 2 | 21093 ± 31 | 23273 ± 1180 |
| F220C | Class 7[1] | Non-Class 2 | 19781 ± 348 | 22066 ± 605 |
| R252P | Unclassified | Non-Class 2 | 18510 ± 536 | 20364 ± 182 |
| del I255 | Unclassified | Severe Class 2 | 1936 ± 36 | 2844 ± 225 |
| del C264 | Unclassified | Severe Class 2 | 1994 ± 106 | 2291 ± 53 |
| G284S | Unclassified | Moderate Class 2 | 13474 ± 730 | 14941 ± 2034 |
| T289P | Unclassified | Moderate Class 2 | 2357 ± 197 | 4688 ± 301 |
| A295V | Class 6[1] | Non-Class 2 | 21819 ± 280 | 22891 ± 1079 |
| S297R | Unclassified | Severe Class 2 | 1638 ± 41 | 2208 ± 185 |
| A298D | Unclassified | Non-Class 2 | 17093 ± 306 | 20131 ± 2116 |
| K311E | Unclassified | Non-Class 2 | 19549 ± 358 | 21928 ± 1173 |
| T320N | Unclassified | Non-Class 2 | 20522 ± 116 | 22270 ± 1060 |
| L328P | Class 1[1] | Non-Class 2 | 22042 ± 342 | 24896 ± 974 |
| A333V | Unclassified | Moderate Class 2 | 12407 ± 392 | 14555 ± 433 |
| E341K | Class 2[1] | Non-Class 2 | 16622 ± 95 | 17258 ± 2084 |
| T342M | Class 1[1] | Non-Class 2 | 24656 ± 401 | 28184 ± 1242 |
| S343C | Unclassified | Non-Class 2 | 19372 ± 323 | 21627 ± 993 |
| Q344P | Class 1[1] | Non-Class 2 | 17471 ± 257 | 18834 ± 1694 |
| Q344R | Class 1[1] | Non-Class 2 | 19609 ± 277 | 22090 ± 1221 |
| V345M | Class 1[1] | Non-Class 2 | 21159 ± 313 | 23618 ± 1023 |
| V345L | Class 1[1] | Non-Class 2 | 20578 ± 243 | 23057 ± 1079 |
| A346P | Class 1[1] | Non-Class 2 | 20301 ± 335 | 22845 ± 1338 |
| P347Q | Class 1[1] | Non-Class 2 | 20697 ± 270 | 23161 ± 1184 |
| P347R | Class 1[1] | Non-Class 2 | 19849 ± 272 | 22682 ± 1304 |
| P347L | Class 1[1] | Non-Class 2 | 22253 ± 148 | 24541 ± 869 |
| P347A | Class 1[1] | Non-Class 2 | 18548 ± 679 | 21170 ± 753 |
| P347S | Class 1[1] | Non-Class 2 | 20830 ± 291 | 23864 ± 986 |


[1] Athanasiou, D., Aguila, M., Bellingham, J., Li, W., McCulley, C., Reeves, P. J., and Cheetham, M. E. (2018) The molecular and cellular basis of rhodopsin retinitis pigmentosa reveals potential strategies for therapy, *Progress in Retinal and Eye Research 62*, 1-23.

[2] Kaushal, S., and Khorana, H. G. (1994) Structure and function in rhodopsin. 7. Point mutations associated with autosomal dominant retinitis pigmentosa, *Biochemistry 33*, 6121-6128.

[3] Sung, C. H., Schneider, B. G., Agarwal, N., Papermaster, D. S., and Nathans, J. (1991) Functional heterogeneity of mutant rhodopsins responsible for autosomal dominant retinitis pigmentosa, *Proceedings of the National Academy of Sciences 88*, 8840.

[4] Wan, A., Place, E., Pierce, E. A., and Comander, J. (2019) Characterizing variants of unknown significance in rhodopsin: A functional genomics approach, *Human Mutation 40*, 1127-1144.
